# An inducible tethered function assay reveals differential regulatory effects for the RNA-binding protein YBX3 and its domains

**DOI:** 10.1101/2025.07.02.662812

**Authors:** William Skipper, Darshan D. Mehta, Justin Adler, Nathan Rose, Isabelle Johnson, Zachary J. Bressman, Michael D. Sheets, Amy Cooke

## Abstract

RNA-binding protein (RBP) regulation is widespread across biology from development to learning and memory. Often RBPs contain multiple modular domains, which contribute to distinct RNA-binding activity or interactions necessary for regulation. However, assays to determine specific regulatory activity of individual domains are limited. The tethered function assay (TFA) provides a direct method to assess the functional activity of RBPs and their domains. The assay consists of two key components: (i) a reporter plasmid that encodes an mRNA, such as luciferase or GFP, engineered with high-affinity binding sites for an exogenous RNA-binding factor, and (ii) an effector construct that expresses a chimera of the RBP fused to the RNA-binding factor. Co-transfection of these constructs allows for measurement of reporter activity as a quantitative readout for RBP regulatory function. We developed an inducible TFA (iTFA) system through generation of a stable inducible mammalian cell line to express a reporter mRNA encoding nanoluciferase (nLuc) with six high-affinity MS2 binding sites within its 3’ untranslated region. These cells can be transfected with a single plasmid that expresses an RBP fused to the MS2 coat protein. This approach enhances efficiency and reproducibility through reduction of transfection burden to a single plasmid and simplified normalization. We used this platform to dissect the individual and combined regulatory effects of YBX3 domains, a modular RBP with differential regulatory activity. The iTFA system provides a streamlined, tunable platform for functional analysis of RBPs that facilitates rapid interrogation of RBP or domain-specific activities in mammalian cells.

## Introduction

Post-transcriptional control permeates biological processes, and RNA-binding proteins (RBP) are major contributors of this control (Bronicki & Jasmin, 2013; Cooke et al., 2019; Corbett, 2018; Gerstberger et al., 2014; Halbeisen et al., 2008; Van Nostrand et al., 2020). Disruption of RBP activity or levels has been implicated in numerous diseases including various cancers, obesity and neurological disorders, which exemplifies their importance and impact on human health (Babic et al., 2004; Crawford Parks et al., 2017; Dai et al., 2015; Tay & Richter, 2001; Xu et al., 2017). Recent advances in system-wide techniques have enabled the global identification of RNAs bound by a single RBP, and the comprehensive effects of protein and mRNA levels upon the RBP perturbation (Van Nostrand et al., 2020). These approaches have revealed a clear feature of RBP control: a single RBP can bind and regulate hundreds of RNAs, yet not all transcripts are regulated equally (Cooke et al., 2019; Van Nostrand et al., 2020). An RBP can perform multiple regulatory activities , which vary among cell types and different RNA targets within a cell. For example, YBX3 represses translation of some targets, while it stabilizes other RNAs it binds to increase protein synthesis, and these targets can vary between species (Awad et al., 2024; Cooke et al., 2019; Van Nostrand et al., 2020). A key challenge, therefore, is to identify features of the RBP that allow it to differentially regulate its targets.

RBPs are often composed of multiple modular domains, which allow a single protein to carry out distinct RNA-binding and protein-protein interaction activities, and enables a wide range of RNA specificity and regulatory control (Corley et al., 2020; Fang et al., 2024; Lunde et al., 2007). These domains may function independently or cooperatively to influence which RNAs or proteins the RBP binds, as well as the regulatory outcomes of these interactions (Corley et al., 2020; Fang et al., 2024). For example, in muscleblind-like (MBNL) proteins the unstructured C-terminal domain anchors MBNL1 to membranes, whereas the zinc finger (ZnF) domains mediate both RNA-binding specificity and kinesin-association, independently of RNA-binding (Hildebrandt et al., 2023). Each of these modular activities is needed for proper RNA localization, which underscores the critical role of domain modularity in RBP function (Hildebrandt et al., 2023). Similarly, vertebrate YBX proteins are characterized by their tripartite modular domain organization: an N-terminal domain rich in alanine and proline, a central cold shock domain (CSD) and a variable C-terminal domain (CTD) that contains alternating acidic and basic patches (Kleene, 2018; Mordovkina et al., 2020). Both the CSD and CTD have RNA-binding activity, and the actions of these domains may be critical for YBX proteins to control the mRNAs they bind (Kleene, 2018; Mordovkina et al., 2020). While it is unclear if these domains act independently or cooperatively to bind and regulate specific RNAs, evidence suggests that the CTD promotes the RNA-binding affinity of the CSD (Izumi et al., 2001).

The tethered function assay (TFA) is an ideal system to investigate how RBPs or their individual domains contribute to regulation (Bos et al., 2016; Coller & Wickens, 2007). The conventional TFA comprises two components: (i) a reporter plasmid encoding an mRNA, such as luciferase or GFP, engineered with high-affinity binding sites for an exogenous RNA-binding factor, such as the MS2 coat protein, and (ii) an effector construct that expresses a chimera of the RBP fused to the RNA-binding factor (Bos et al., 2016; Coller & Wickens, 2007). Following co-transfection, reporter activity is measured to quantify RBP-directed mRNA regulation. However, the conventional approach to the TFA requires substantial optimization to ensure robust effects on the reporter, expression of the RBP chimera and normalization (Bos et al., 2016; Coller & Wickens, 2007). In addition, recent reports suggest that transient transfections can lead to limited cellular resources for effective plasmid expression that may contribute assay variability (Di Blasi et al., 2021; Frei et al., 2020; Jones et al., 2020).

To overcome these limitations, we developed an inducible TFA (iTFA). The key component is a stable HeLa cell line that can be induced to express a reporter mRNA with high-affinity MS2 binding sites (Spitzer et al., 2013). This approach increases efficiency and reproducibility compared to the transient TFA by reducing the transfection burden to a single effector plasmid and ensures uniform reporter expression across the cell population, which simplifies normalization (Figure 1A). The iTFA reporter consists of a nanoluciferase (nLuc) mRNA that contains six MS2 stem loops in the 3′ untranslated region (UTR) that enable specific binding by MS2 coat protein RBP chimeras. Upon transfection with known MS2-fused regulatory protein chimeras, such as CNOT7 or BOLL, the reporter exhibits the expected changes in luciferase activity, which validates the system (Luo et al., 2020). Using this platform, we dissected the individual and combined regulatory effects of the YBX3 domains. This iTFA system provides a streamlined, tunable platform for functional analysis of RBPs, offering precise control of reporter RNA levels and facilitating rapid interrogation of RBP or domain-specific activities in mammalian cells.

**Figure 1.**
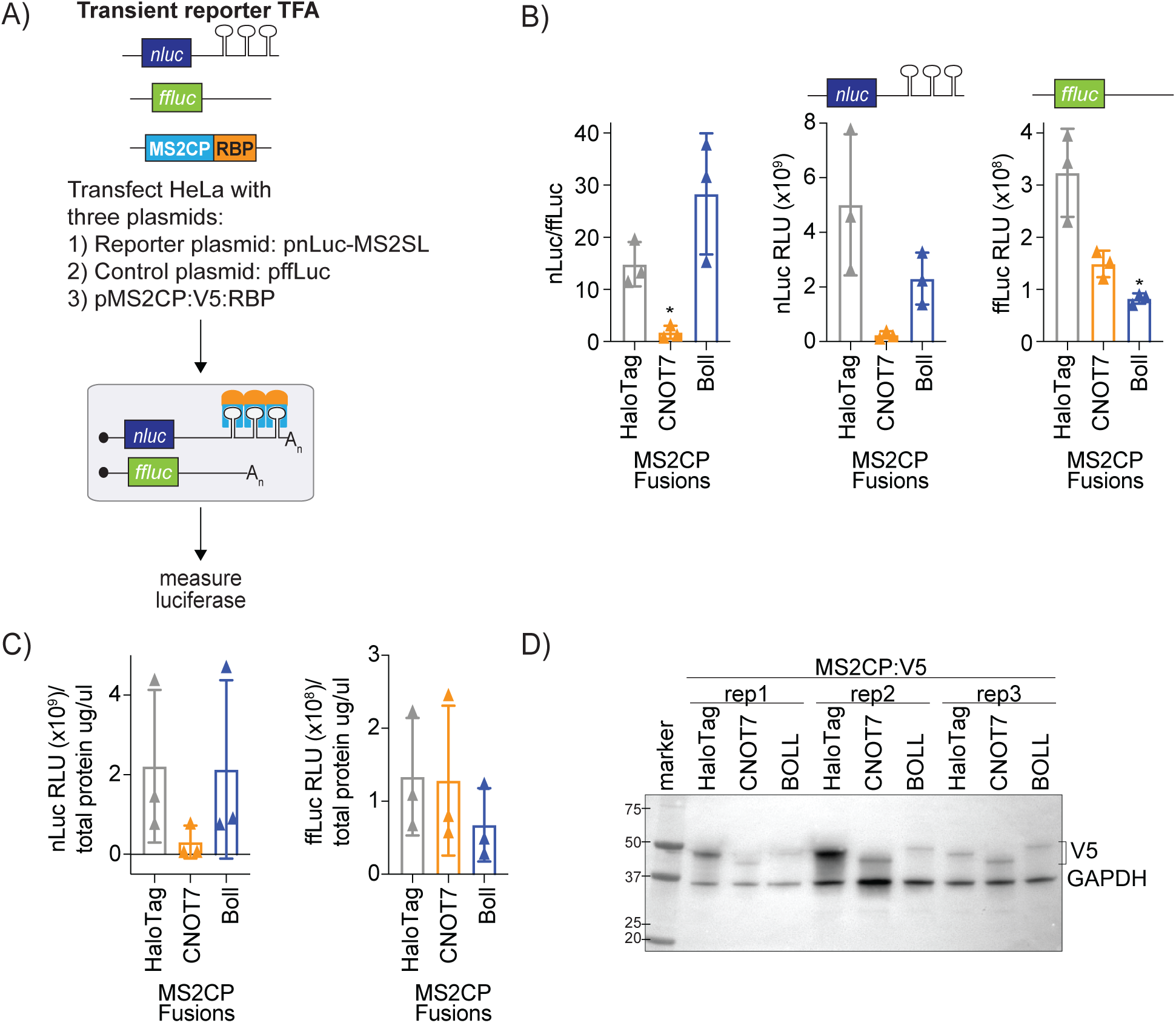
Transient reporter tethered function assay produces variable results. A) Schematic of the established transient reporter tethered function assay (TFA). Plasmids encoding two different reporters (nanoluciferase (nLuc) with MS2 stem loops (SL) within the 3’ UTR and firefly luciferase (ffLuc) lacking MS2 SL) and a plasmid encoding a MS2 coat protein V5 (MS2CP:V5) RNA-binding protein (RBP) chimera are co-transfected into cells. B) Luciferase assay of normalized reporter expression (*left)* nLuc/ffLuc (*middle)* nLuc Relative Light Units (RLU) and (*right)* ffLuc RLU with the indicated MS2CP:V5:RBP effector chimeras, CNOT7 or BOLL, and the HaloTag control. All luciferase data are displayed as single points, mean +/- standard deviation, n=3. C) (*left)* nLuc RLU and (*right)* ffLuc RLU normalized to total protein concentration (µg/µl) of the indicated MS2CP:V5:RBP effector chimeras, CNOT7 or BOLL, and the HaloTag control. All luciferase data are displayed as single points, mean +/- standard deviation, n=3. D) Immunoblot analysis of all three replicates of MS2CP:V5 chimeras, HaloTag, CNOT7 and BOLL, with GAPDH as a loading control. Molecular weight markers are indicated.

## Results

### The transient reporter tethered function assay produces variable results

The conventional TFA requires transfection of three plasmids: two reporter plasmids and one plasmid expressing the MS2 coat protein (MS2CP):RBP chimera (Figure 1A, schematic). To assess the variability and consistency of this system, we tethered known translational regulators with well-characterized effects. When tethered, CNOT7 (CCR4-NOT transcription complex subunit 7) and BOLL (Boule homolog, RNA-binding protein) are known to repress and enhance reporter expression, respectively (Cooke et al., 2010; Enwerem et al., 2021; Luo et al., 2020; Zhang et al., 2013). Therefore, we generated plasmids encoding MS2CP:V5:CNOT7 and MS2CP:V5:BOLL fusion proteins. These were transfected into HeLa cells that were then co-transfected 24 hours later with the nanoluciferase (nLuc) reporter containing six MS2 stem loops (SL) in the 3’ untranslated region (UTR) and the firefly (ffLuc) control plasmid. As shown in Figure 1B, the raw luminescence signals from the nLuc (middle graph) and ffLuc (right graph) reporters display variability as indicated by the large error bars across replicates and constructs, which obscures the expected regulatory effects of the tethered proteins. Normalization (nLuc/ffLuc) partially mitigates this variability, as it causes the repressive effects of CNOT7 to become statistically significant; however, the BOLL construct fails to produce a significant increase in normalized reporter activity as expected (Figure 1B, left graph). One possibility for the variation of the luciferase signal could be a difference in cell number between conditions; however, normalization of the raw luminescence of either nLuc or ffLuc to total protein concentration as a proxy for cell number does not reduce the variability (Figure 1C). This disconnect between the raw and normalized values, and the lack of a robust BOLL effect even after normalization, likely reflects intrinsic noise of transient expression (Figure 1D) and cellular stress introduced through transfection of three plasmids (Di Blasi et al., 2021; Frei et al., 2020; Jones et al., 2020). This variability and limited sensitivity highlights the need for a more streamlined and reliable system to detect regulatory effects.

### Establishing the inducible tethered function reporter cell line

To address the limitations of a transient TFA, we developed an inducible tethered function assay (iTFA) system to reduce the transfection burden and improve reproducibility. We used the Flp-In T-REx system to generate an isogenic inducible HeLa cell line that stably expresses a nanoluciferase (nLuc) reporter with six MS2 SL in the 3’ UTR, referred to as the iTFA cell line. Genomic integration of the reporter allows normalization to cell count or total protein rather than a co-transfected control plasmid. In this streamlined setup, only the MS2CP:V5:RBP chimera needs to be transfected into these cells followed by induction with doxycycline (Dox) (Figure 2A, schematic). To test that the nLuc:MS2SL reporter was integrated and inducible, cells were treated with varying Dox concentrations. The nLuc levels measured were ∼100-fold greater than the uninduced cells, with strong induction observed across all Dox concentrations (Figure 2B). These results demonstrate that the iTFA cell line expresses the nLuc:MS2SL reporter RNA upon Dox- induction.

**Figure 2.**
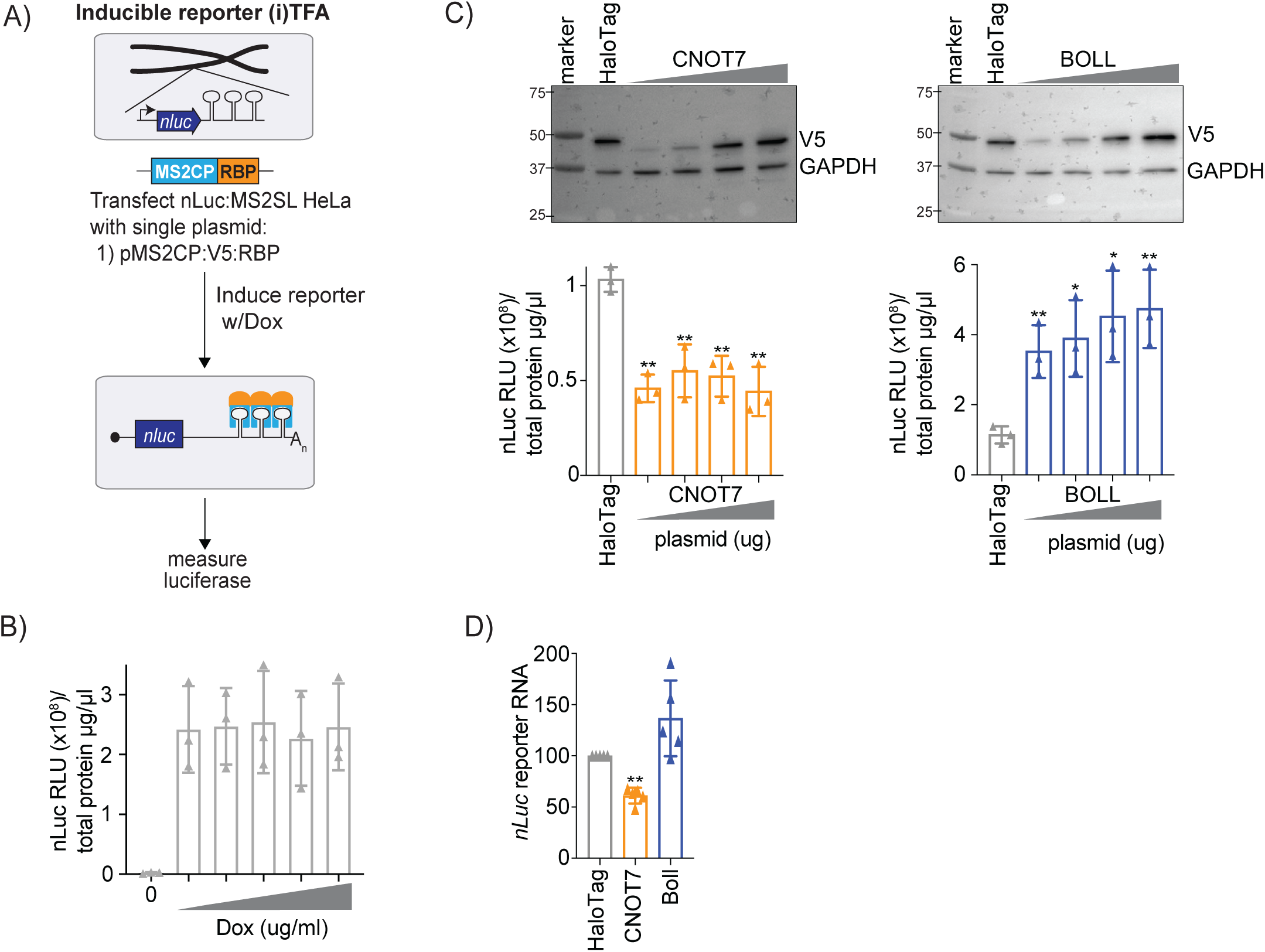
Establishing the inducible tethered function assay with known translational regulators. A) Schematic of the newly established inducible tethered function assay (iTFA). The iTFA uses a HeLa cell line with the nLuc reporter integrated in the genome using the Flp-In T- REx system. Plasmids encoding MS2CP:V5:RBP are transfected into these cells and reporter expression is induced with doxycycline (Dox) ∼16 hours before measuring luciferase. B) Luciferase assay of normalized nLuc reporter expression (nLuc RLU/total protein µg/µl) with the indicated Dox concentrations (0, 50, 100, 200, 500 and 1000 ng/ml) ∼16 hours post induction. All nLuc data are displayed as single points, mean +/- standard deviation, n=3. C) iTFA with MS2 coat protein (MS2CP:V5) effector chimeras, CNOT7 or BOLL, and the HaloTag control. Briefly, iTFA cells were transfected with the indicated plasmids encoding MS2CP:V5 RBP using different amounts of plasmids (0.4µg, 0.8µg, 1.6µg and 3.2µg) followed by doxycycline induction of the nLuc reporter RNA with 6 MS2 stem loops in the the 3’ UTR ∼16 hours before harvesting. (*Top*) Representative immunoblot analysis of MS2 coat protein V5 (MS2CP:V5) effector, CNOT7 or BOLL, with GAPDH as a loading control. Molecular weight markers indicated. (*Bottom*) Normalized nLuc reporter measurements for tethered CNOT7 or BOLL. Total protein concentration from each cell lysate was used to normalize nLuc values across samples. All nLuc data are displayed as single points, mean +/- standard deviation. * = p < 0.05, ** = p < 0.01, and *** = p < 0.001 with Student’s t-test, n=3. D) Reverse transcription quantitative polymerase chain reaction (RT-qPCR) of relative nLuc reporter levels from iTFA with the indicated MS2CP:V5 fusion protein. * = p < 0.05 with paired Student’s t-test, n=3. RT-qPCR values normalized to *GUSB* mRNA levels.

### The iTFA system reflects activities of known translational regulators

Next, we sought to determine whether the iTFA system could replicate the effects of the known translational regulators, CNOT7 and BOLL. We titrated the concentration of transfected plasmids encoding MS2CP:V5:CNOT7 and MS2CP:V5:BOLL to optimize chimera expression (Figure 2C, top blots). After Dox induction of the reporter, we measured nLuc activity to assess the effects of tethered MS2CP:V5:CNOT7 or MS2CP:V5:BOLL. As expected, MS2CP:V5:CNOT7 significantly reduced nLuc levels, while MS2CP:V5:BOLL significantly increased reporter activity compared to the MS2CP:V5:HaloTag control (Figure 2C, bottom). We next analyzed nLuc reporter RNA levels as CNOT7 is known to reduce RNA levels while BOLL has little to no effect on the levels of reporter RNA when tethered (Luo et al., 2020). MS2CP:V5:CNOT7 significantly reduced nLuc reporter RNA, while MS2CP:V5:BOLL displayed no effect (Figure 2D). These results show that the iTFA system reproduces both repressive and stimulatory regulatory effects of the tethered proteins, mirrored in reporter RNA levels, and reduces the number of transfected plasmids.

### The iTFA system can be used to assess RBP domain activity

One major advantage of the TFA system is direct analysis of the regulatory effects of individual RBP domains independent of the context of the full-length RBP (Bos et al., 2016; Coller & Wickens, 2007). Therefore, we tested if the iTFA system can replicate RBP domain activity using BICC1 (Bicaudal C Homolog 1) and its domains, as have been previously tested in a transient TFA (Zhang et al., 2013). As expected, the full-length (FL) MS2CP:V5:BICC1 and the BICC1 C-terminal domain (CTD) chimeras significantly decreased nLuc activity when tethered in the iTFA, whereas the MS2CP:V5 N-terminal domain (NTD) chimera did not alter nLuc activity compared to the MS2CP:V5:HaloTag control (Figure 3A). MS2CP:V5:CNOT7 and MS2CP:V5:BOLL were used as positive controls to ensure that the iTFA assay was functional. The BICC1 domain constructs expressed at higher levels than the MS2CP:V5:BICC1 construct (Figure 3B), which is likely due to the size of the plasmids transfected, which can cause differences in transfection efficiency (Kreiss et al., 1999). The BICC1 and domains combined with the MS2CP:V5:CNOT7 and MS2CP:V5:BOLL results validate that the iTFA system recapitulates previous results observed with the transient TFA but with a more effective, simplified experimental approach.

**Figure 3.**
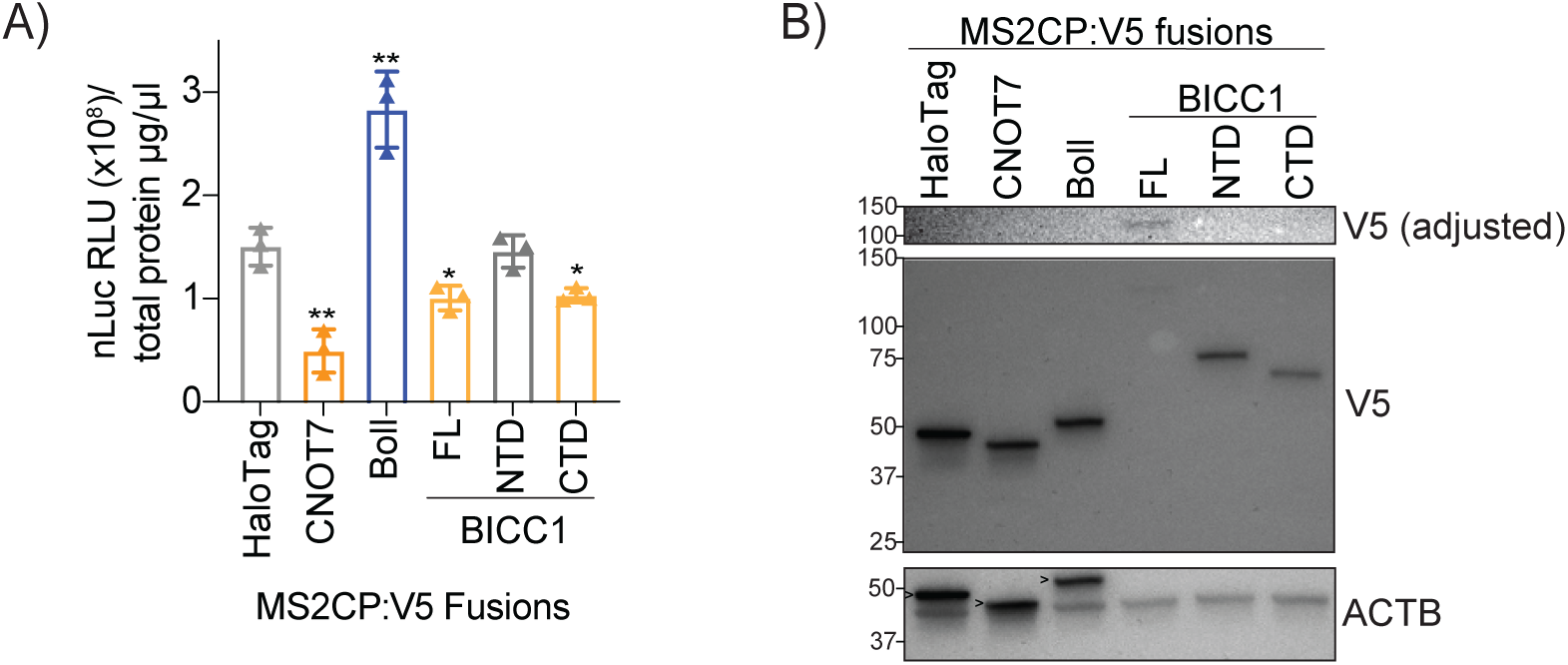
BIC-C and its domains exert predictable effects on nLuc activity when analyzed using the iTFA. A) iTFA with plasmids encoding MS2CP:V5 chimera controls HaloTag, CNOT7 and BOLL along with BIC-C full-length (FL) or individual N-terminal domain (NTD) or C-terminal domain (CTD). Normalized nLuc reporter expression (nLuc RLU/total protein µg/µl) plotted. All nLuc data are displayed as single points, mean +/- standard deviation. * = p < 0.05 and ** = p < 0.01 with Student’s t-test, n=3. B) Representative immunoblot analysis of MS2CP:V5 chimera expression in iTFA along with actin loading control (ACTB). Protein molecular weights indicated and > indicates leftover V5 antibody signal in ACTB blot. The V5 adjusted image has had the contrast increased on the V5 image to show the expression of FL BIC-C chimera.

### Full-length YBX3 increases luciferase activity when tethered

The YBX RBP family is distinguished by its tripartite domain organization (Figure 4A) (Mordovkina et al., 2020). In particular, YBX3 has been shown to mediate post-transcriptional control mechanisms that can either inhibit protein synthesis of target mRNAs or promote mRNA stability to increase protein levels of its targets (Cooke et al., 2019). We set out to determine if the modular domain organization of YBX3 dictates its post-transcriptional regulatory functions using the iTFA system. To do this, we generated plasmids encoding full-length and individual domain MS2CP:V5:YBX3 chimeras to test in the iTFA cell line (Figure 4A, schematics). These were transfected into iTFA cells and verified all MS2CP:V5:YBX3 chimeras were expressed (Figure 4B). As with the BOLL chimera, the full-length MS2CP:V5:YBX3 chimera significantly increased nLuc activity when tethered, while the YBX3 individual domains (NTD, cold shock domain (CSD) and CTD) had little to no effect on the nLuc activity when compared to the control MS2CP:V5:HaloTag (Figure 4C). At the RNA level, the full-length MS2CP:V5:YBX3 and individual domains had little to no effect similar to MS2CP:V5:HaloTag (Figure 4D). The positive (MS2CP:V5:BOLL) and negative (MS2CP:V5:CNOT7) effector controls exhibited the expected expression and activities (Figure 4B-D). This data indicates that when tethered, full-length YBX3 activates nLuc reporter expression independent of reporter RNA levels, consistent with translational activation, while the individual domains are inert.

**Figure 4.**
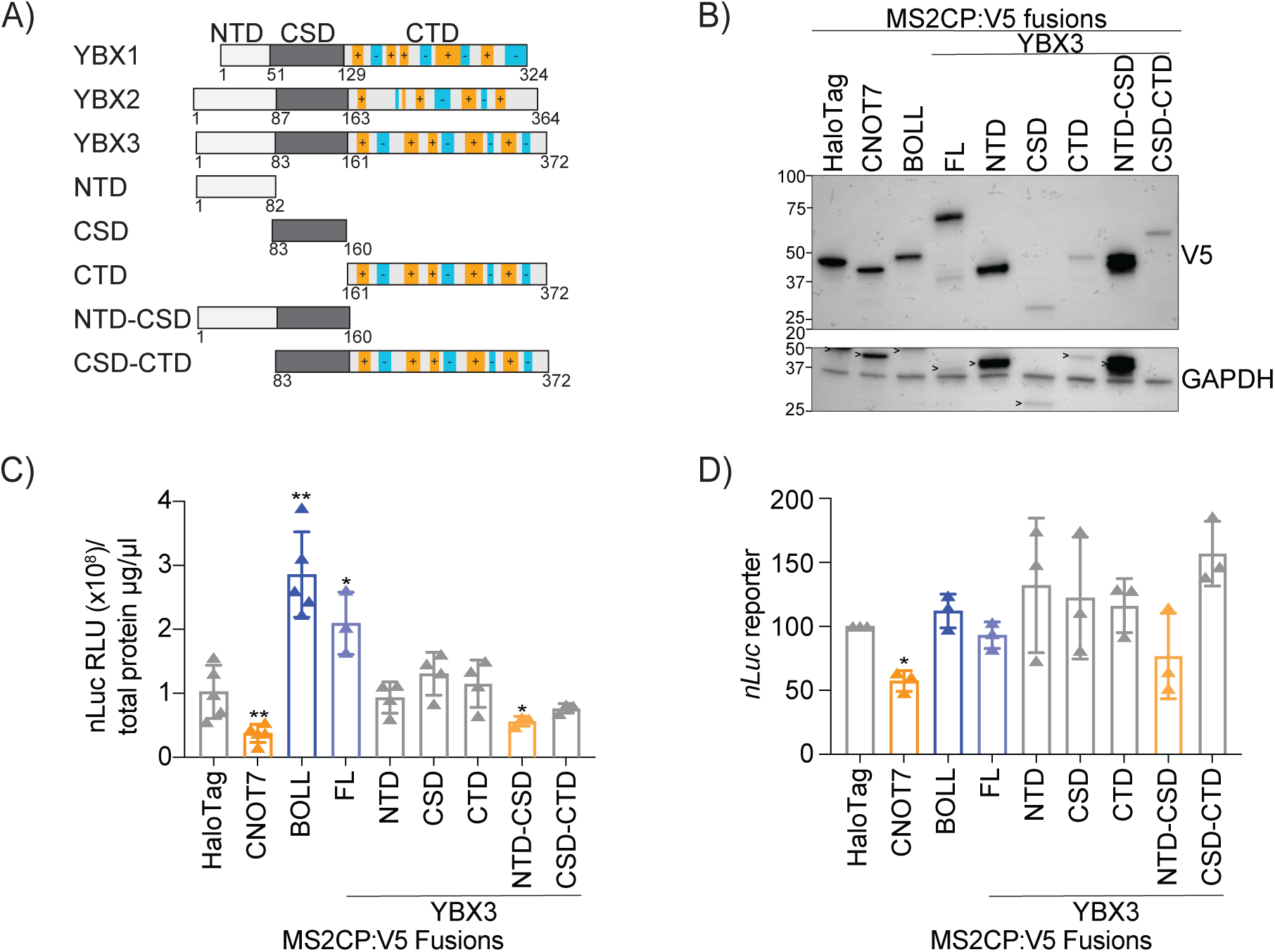
Tethered YBX3 and its domains exhibit differential effects on nLuc activity. A) Schematic of YBX domain organization for the vertebrate YBX proteins (YBX1, YBX2 and YBX3) and YBX3 individual and domain combinations: N-terminal domain (NTD), cold shock domain (CSD) or C-terminal domain (CTD). B) Representative immunoblot analysis of MS2CP:V5 chimera expression of HaloTag, CNOT7, BOLL and YBX3 constructs depicted in (A) along with GAPDH loading control. Protein molecular weights indicated and > indicates leftover V5 antibody signal in GAPDH blot. C) Normalized nLuc reporter expression (nLuc RLU/total protein µg/µl) plotted. All nLuc data are displayed as single points, mean +/- standard deviation. * = p < 0.05, and ** = p < 0.01 with Student’s t-test, n<3. D) RT-qPCR analysis of relative nLuc reporter levels from iTFA with the indicated MS2CP:V5 fusion protein. * = p < 0.05 with paired Student’s t-test, n=3. RT-qPCR values normalized to *GUSB* mRNA levels.

### YBX3 lacking the CTD exhibits a reduction in luciferase activity when tethered

As the individual YBX3 domains exhibited modest effects on nLuc activity when tethered, we next tested if domain combinations could elicit regulatory effects using the iTFA. We generated the different possible domain combination chimeras, (Figure 4A, schematic) then transfected these plasmids into the iTFA reporter cell line. Similar to the reduction in nLuc activity with tethered MS2CP:V5:CNOT7, the domain combination that contained the YBX3 NTD and CSD (MS2CP:V5:NTD-CSD) decreased the amount of nLuc activity, while the CSD-CTD domain combination had little to no effect when tethered (Figure 4C). Analysis of reporter RNA levels showed that the neither combination reduced RNA abundance (Figure 4D). All domain combination chimeras were expressed and the controls elicited the expected effects (Figure 4B- D). These combined data suggest that when tethered the YBX3 NTD works in combination with the CSD to translationally repress expression, while the full-length protein translationally activates nLuc expression.

## Discussion

Here, we developed and validated an inducible tethered function assay (iTFA) as an improved alternative to the conventional transient TFA, which has increased variability likely due to the high transfection burden (Figure 1). The iTFA system leverages a doxycycline-inducible reporter stably integrated into the genome (a HeLa cell line), which ensures consistent reporter expression and reduces the number of transfected plasmids to a single MS2CP:V5:RBP chimera. This design improves reproducibility, simplifies normalization, and reduces resource competition effects associated with transient transfections (Di Blasi et al., 2021; Frei et al., 2020; Jones et al., 2020). We validated the iTFA with known translational regulators, CNOT7 and BOLL, which produced expected activation or repression on nLuc activity and on reporter RNA abundance (Figure 2) (Cooke et al., 2010; Enwerem et al., 2021; Luo et al., 2020). In addition, we used BICC1 and its domains to demonstrate that iTFA reproduces known domain-specific effects, consistent with previous transient TFA effects (Figure 3) (Zhang et al., 2013). Finally, we used the iTFA system to dissect the regulatory contributions of YBX3 and its domains, which revealed that full-length YBX3 activates translation independent of increased reporter RNA levels, while the NTD- CSD domain combination repressed translation without affecting reporter RNA abundance (Figure 4). These results establish iTFA as a new, versatile and reliable system to analyze RNP and domain-specific functions in mammalian cells.

Our analysis of YBX3 reveals how its modular domains contribute to distinct regulatory activities. Tethered full-length YBX3 increased nLuc activity independent of changes to reporter RNA levels, which indicates that the protein activates translation directly. Previous reports have shown YBX3 stabilizes mRNAs that in turn increases protein synthesis (Cooke et al., 2019; Nie et al., 2012). Our results here suggest that YBX3 can also stimulate translation independent of mRNA stability. In contrast, the tethered NTD-CSD domain combination reduced luciferase activity and did not alter reporter RNA levels, which suggests this domain combination represses translation. Neither domain alone affected nLuc protein or RNA expression, which implies that YBX3 requires cooperation between regions to regulate expression. These results align with prior studies that show YBX proteins exert dual regulatory effects through complex interactions between domains (Izumi et al., 2001; Kleene, 2018; Lindquist & Mertens, 2018; Mordovkina et al., 2020). Interestingly, when tethered in zebrafish embryos, full-length YBX1 repressed translation (Sun et al., 2018), which may reflect a context-dependent regulation, such as differences in cell type, species, or functional variation between proteins. A recent study further demonstrates YBX3 differentially regulates amino acid transporter mRNAs in the species-dependent manner (Awad et al., 2024). Future studies could investigate whether domain-specific effects result from distinct protein-protein interactions or changes in RNA-binding conformation, and how these differences contribute to functional divergence across the YBX family.

The iTFA system offers a streamlined, sensitive platform to interrogate the regulatory functions of RBPs and their domains. It requires the construction and transfection of just one type of plasmid, the MS2CP:V5:RBP chimera, which reduces optimization of the conventional TFA and variability tied to resource competition with co-transfections (Di Blasi et al., 2021; Frei et al., 2020; Jones et al., 2020). While we developed this in a Flp-In T-REx HeLa cell line, any Flp-In T-REx cell could be used to generate an iTFA reporter line, e.g., Flp-In T-REx HEK293T (Spitzer et al., 2013), which expands the utility of this method. Given the iTFA’s ability to resolve differential regulatory effects of complete RBPs and their domains, this system could be used to dissect other complex RBPs. For example, the Pumilio proteins PUM1 and PUM2 regulate their mRNA targets through diverse mechanisms (Bohn et al., 2018; Enwerem et al., 2021; Kaye et al., 2009; Uyhazi et al., 2020; Van Etten et al., 2012). While their binding structures and target motifs are similar, the two often display opposing regulatory effects (Uyhazi et al., 2020). The iTFA could determine the domains responsible to activate or repress by expressing individual or combined regions of PUM1 and PUM2 as MS2CP chimeras. A conserved amino acid stretch in the NTD of PUM1 and PUM2 mediates translational repression through the recruitment of CNOT7 (Enwerem et al., 2021), but the region for activation remains undefined. The iTFA system could map the region(s) needed for activation and clarify how each PUM protein contributes to distinct post-transcriptional regulatory outcomes. Similarly, the protein G3BP1 represses specific mRNAs in nerve cells and inhibiting its activity has been shown to improve regeneration in damaged nerve tissue (Sahoo et al., 2018; Sidibe et al., 2021). By identifying which region of the protein is responsible for this regulation with the iTFA, a therapeutic target could be revealed that may be simpler and less detrimental to disable than targeting the entire protein.

## Acknowledgements

We would like to thank the past and present members of the entire Cooke lab for critical discussions, helpful feedback and general support. We are grateful to the members of the Haverford Biology support staff that offered assistance throughout this study, in particular, Nicole Cunningham, Zak Kerrigan, Ebonee Jackson and Joanne Brown. We would also like to thank Geoffrey Hutinet for helpful suggestions on figures. We would also like to thank Elena Dobrikova and Matthias Gromeier (at Duke University Medical Center) for the provision of the HeLa Flp-In T-REx cell line needed for all inducible stable cell lines produced in this study. Finally, A.C. thanks the NIH NIGMS (1R15GM146205-01) for grant support that funded this work. This research was also supported by NICHD (R01HD091921) and NIGMS (R01GM152615) grants to MDS.

## Materials and Methods

### Cell culture and Generation of iTFA Inducible Cell Line

Adherent HeLa cells (ATCC Cat# CCL-2, RRID:CVCL_0030) were grown at 37 °C and 5% CO_2_ in DMEM, supplemented with 10% FBS Premium Plus (Gibco), 1% Penicillin/Streptomycin (Gibco) and 1% L-glutamine (Gibco). To generate the iTFA Doxycycline-Inducible reporter cell line, the pCDNAFRT TO NanoLuc+6xMS2SL was transfected into the HeLa Flp-In T-REx cell line then selected as described in the manufacturer’s protocol (Flp-In T-REx, Invitrogen). To confirm nLuc integration, a luciferase assay was performed according to the manufacturer’s recommendations (Nano-Glo Luciferase Assay System, Promega, cat# N1110). Luciferase activity was measured using a Promega GloMax 20/20 Luminometer, using the preset Promega protocol Luc-0-Inj.

### Plasmids Generated

#### Reporter plasmids

Nanoluciferase (nLuc) is a luciferase enzyme derived from the deep-sea shrimp *Oplophorus gracilirostris* (England et al., 2016). The nLuc plasmid (pCDNAFRT TO NanoLucMS2x6) used in the creation of the iTFA cell line and the transient TFA was generated in Michael Sheets’ laboratory at the University of Wisconsin-Madison. pGL13.4 expresses firefly luciferase lacking the MS2 stem loops (Promega).

#### MS2-RBP Fusion Proteins

All the MS2-RBPs used were expressed from the vector backbone pFN21a. The control plasmid, pFN21A:Halo:MS2CP:V5 was used as a negative control (Van Etten et al., 2012), which was a gift from the laboratory of Aaron Goldstrohm at the University of Minnesota. The HaloTag was removed by NheI/XbaI restriction digest and the MS2CP:V5 was reinserted via Gibson Assembly in the NheI/XbaI restriction sites generating pFN21a:MS2CP:V5 and all RBPs were cloned into this plasmid via Gibson assembly into the NheI/XbaI restriction sites except for the YBX3 chimeras that contained the NTD. For these, YBX3 was cloned by General Biosystems into pFN21A:Halo:MS2CP:V5. To generate the MS2CP:V5 plasmids for FL, NTD, and NTD:CSD, the HaloTag was removed with NheI/PvuI restriction digest and a small oligo inserted to replace the T7 promoter and Kozak sequnece. The pFN21a:Halo:MS2CP:V5:CNOT7 (Van Etten et al., 2012) (from the Goldstrohm lab) and egfp- BicC-HA-V5turbo (from the Sheets lab at the University of Wisconsin-Madison) were used to amplify these inserts. The primers used for this Gibson assembly are listed in Table 2 below.

**Table 1:** Plasmid mass for transfection.

| plasmid | mg transfected |
| --- | --- |
| pFN21a:MS2CP:V5 | 1.6 mg |
| pFN21a:MS2CP:V5:CNOT7 | 1.6 mg |
| pFN21a:MS2CP:V5:BOLL | 1.6 mg |
| pFN21a:MS2CP:V5:BICC1 | 4.2 mg |
| pFN21a:MS2CP:V5:BICC1 NTD | 2.5 mg |
| pFN21a:MS2CP:V5:BICC1 CTD | 2.3 mg |
| pFN21a:MS2CP:V5:YBX3 | 2.0 mg |
| pFN21a:MS2CP:V5:YBX3 NTD | 1.0 mg |
| pFN21a:MS2CP:V5:YBX3 CSD | 1.0 mg |
| pFN21a:MS2CP:V5:YBX3 CTD | 1.3 mg |
| pFN21a:MS2CP:V5:YBX3 NTD:CSD | 1.2 mg |
| pFN21a:MS2CP:V5:YBX3 CSD:CTD | 1.6 mg |

**Table 2:**
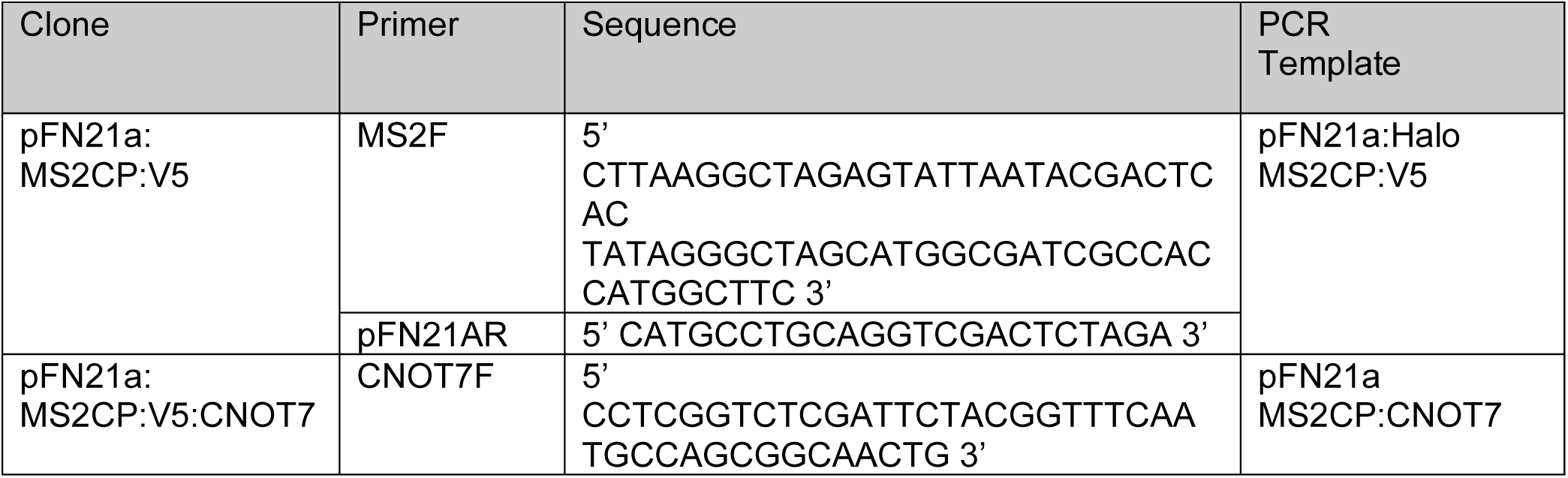
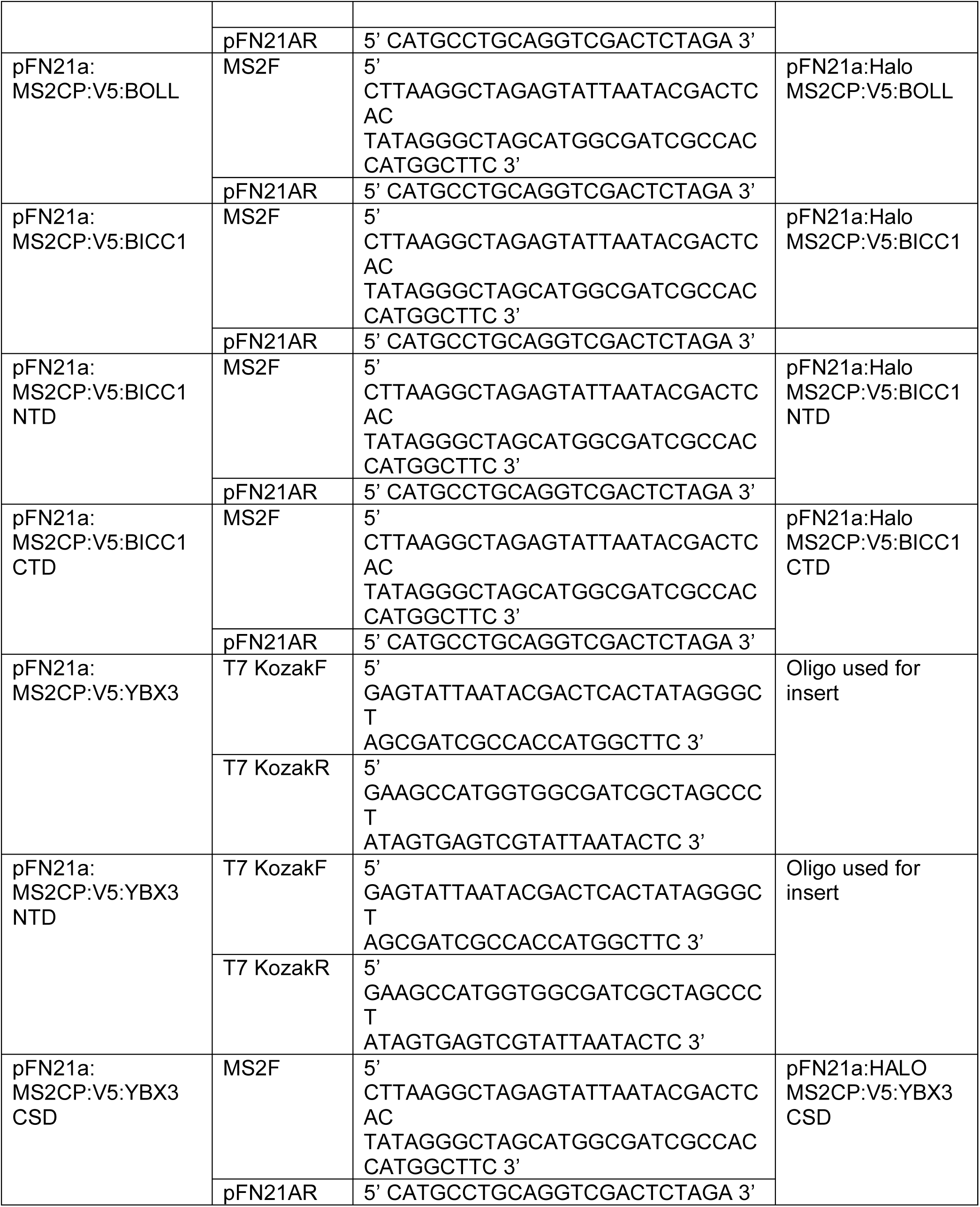
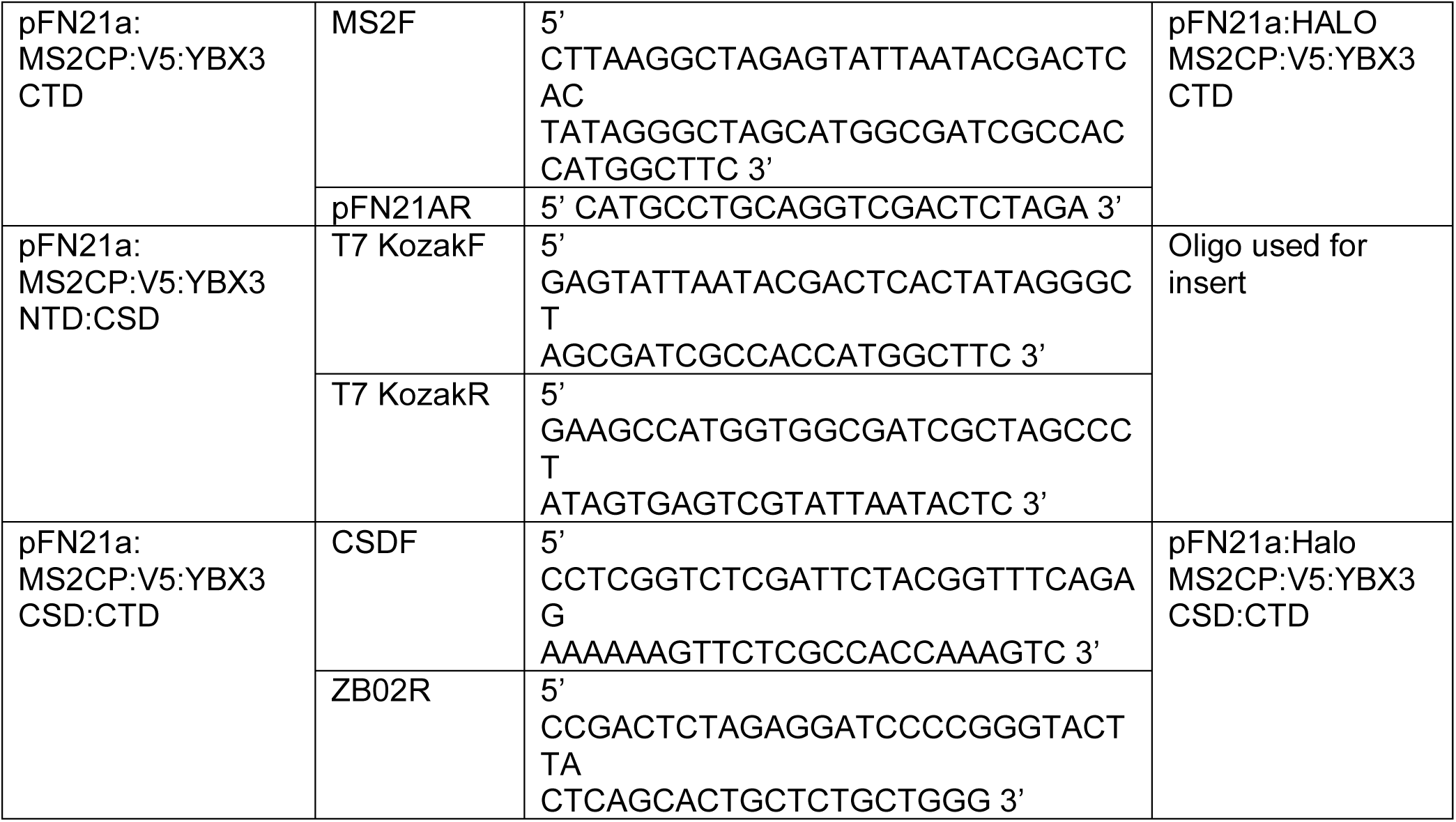
Primers for Gibson Assembly.

#### Transient TFA and iTFA Transfection Procedure

HeLa cells or iTFA reporter cells were grown until ∼70-80% confluent, trypsinized, counted (Biorad TC20) and plated in 6-well plates to achieve ∼60-70% confluence 24 hours post-seeding. The cells were then transfected with the µgindicated of pFN21a:MS2CP:V5 chimeras in Table 1 below using lipofectamine 2000 (Invitrogen) following manufacturer’s recommendation. The cells were grown for 24 hours at 37 °C and 5% CO_2_ to allow expression of the MS2CP:V5 chimeras. iTFA cells were then induced with 50 ng/ml doxycycline to induce reporter expression, while the transient TFA cells were co-transfected with 500ng pCDNAFRT TO NanoLuc+6xMS2SL and 1µgpGL13.4 plasmids. After 24 hours, cells were washed with 1x PBS and harvested in 1x PLB (Promega) for luciferase, Western blot and RNA analysis.

#### Luciferase Assay

TFA cell lysates were assayed for luciferase expression using either Nano-Glo Luciferase Assay System, Promega, cat# N1110 for iTFA or Nano-Glo Dual-Luciferase Reporter Assay System, Promega, cat# N1610 for the transient TFA following manufacturer’s recommendations. Luciferase activity was measured using a Promega GloMax 20/20 Luminometer, using the preset Promega protocol Luc-0-Inj for the iTFA or Dual-0-Inj for the transient TFA.

### Western Blots

Protein concentration of cell lysates from transient TFA or iTFA were determined using Protein Assay Dye (Bio-Rad, cat# 5000006) following manufacturer’s recommendation with Pharmacia LKB Ultrospec III UV/Vis spectrophotometer. 10-15µgof total protein were run on a 4-15% TGX gel (Biorad) in 1x Laemmli running buffer, transferred to nitrocellulose membranes using the standard turbo transfer mixed molecular weight program on the Trans Blot Turbo System (Biorad), and analyzed using primary antibodies. The primary antibodies used in immunoblotting were anti- V5 tag (Invitrogen, cat# R960-25), 1:10000, anti- -ACTIN clone C4 (Sigma Aldrich, MAB1501), 1:5000, and anti-GAPDH (Proteintech, cat# 60004-1), 1:10000. Secondary antibody used was Goat Anti-Mouse IgG H&L HRP (Abcam, ab205719), 1:10000.

### RNA extraction and RT-qPCR analysis

Total RNA was isolated using GeneJet RNA Purification kit (#K0731, Thermo Fisher) kit following the manufacturer’s recommendations. 250-500 ng total RNA was used to synthesize cDNA using the Superscript III First Strand Synthesis System (#18080051, Thermo Fisher) for quantitative reverse transcription polymerase chain reaction (RT-qPCR). RT-qPCR was performed using the SYBR-Green qPCR Master mix (#4309155, Applied Biosystems) on a QuantStudio3 Real-Time PCR System (Thermo Fisher). Gene expression values were normalized using *GUSB* and are shown as a relative fold change to the value of HaloTag transfected samples using the DDCT method. All experiments were performed in biological triplicate and error bars indicate +/- standard deviation as assayed by the Ct method. A Student’s t-test was used to determine significance in values for HaloTag control versus MS2CP:V5:RBP chimeras. All RT-qPCR primers are listed in the Table 3 below.

**Table 3:**
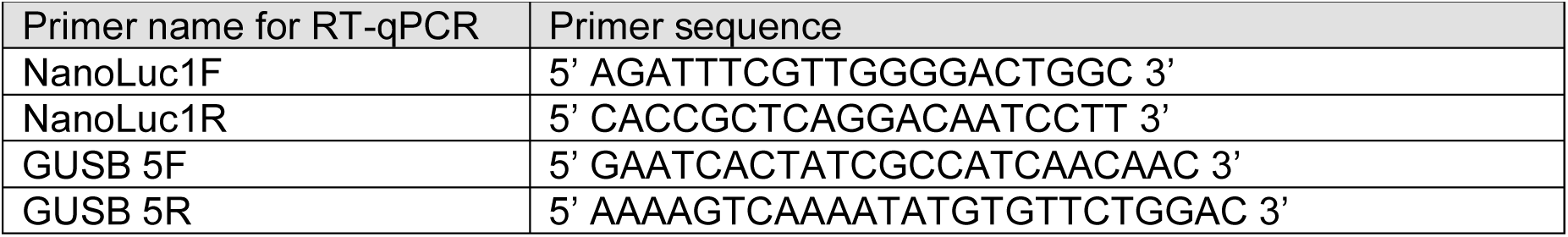
qPCR primers.

## Notes

### Competing Interest Statement

The authors have declared no competing interest.

## References

1. Awad, S., Skipper, W., Vostrejs, W., Ozorowski, K., Min, K., Pfuhler, L., Mehta, D., & Cooke, A. (2024). The YBX3 RNA-binding protein posttranscriptionally controls SLC1A5 mRNA in proliferating and differentiating skeletal muscle cells. Journal of Biological Chemistry, 300(2), 105602. 10.1016/j.jbc.2023.105602

2. Babic, I., Jakymiw, A., & Fujita, D. J. (2004). The RNA binding protein Sam68 is acetylated in tumor cell lines, and its acetylation correlates with enhanced RNA binding activity. Oncogene, 23(21), 3781–3789. 10.1038/sj.onc.1207484

3. Bohn, J. A., Van Etten, J. L., Schagat, T. L., Bowman, B. M., McEachin, R. C., Freddolino, P. L., & Goldstrohm, A. C. (2018). Identification of diverse target RNAs that are functionally regulated by human Pumilio proteins. Nucleic Acids Res, 46(1), 362–386. 10.1093/nar/gkx1120

4. Bos, T. J., Nussbacher, J. K., Aigner, S., & Yeo, G. W. (2016). Tethered Function Assays as Tools to Elucidate the Molecular Roles of RNA-Binding Proteins. Rna Processing: Disease and Genome-Wide Probing, 907, 61–88. 10.1007/978-3-319-29073-7_3

5. Bronicki, L. M., & Jasmin, B. J. (2013). Emerging complexity of the HuD/ELAVl4 gene; implications for neuronal development, function, and dysfunction. RNA, 19(8), 1019–1037. 10.1261/rna.039164.113

6. Coller, J., & Wickens, M. (2007). Tethered function assays: an adaptable approach to study RNA regulatory proteins. Methods Enzymol, 429, 299–321. 10.1016/S0076-6879(07)29014-7

7. Cooke, A., Prigge, A., & Wickens, M. (2010). Translational repression by deadenylases. J Biol Chem, 285(37), 28506–28513. 10.1074/jbc.M110.150763

8. Cooke, A., Schwarzl, T., Huppertz, I., Kramer, G., Mantas, P., Alleaume, A. M., Huber, W., Krijgsveld, J., & Hentze, M. W. (2019). The RNA-Binding Protein YBX3 Controls Amino Acid Levels by Regulating SLC mRNA Abundance. Cell Rep, 27(11), 3097–3106 e3095. 10.1016/j.celrep.2019.05.039

9. Corbett, A. H. (2018). Post-transcriptional regulation of gene expression and human disease. Curr Opin Cell Biol, 52, 96–104. 10.1016/j.ceb.2018.02.011

10. Corley, M., Burns, M. C., & Yeo, G. W. (2020). How RNA-Binding Proteins Interact with RNA: Molecules and Mechanisms. Mol Cell, 78(1), 9–29. 10.1016/j.molcel.2020.03.011

11. Crawford Parks, T. E., Ravel-Chapuis, A., Bondy-Chorney, E., Renaud, J. M., Cote, J., & Jasmin, B. J. (2017). Muscle-specific expression of the RNA-binding protein Staufen1 induces progressive skeletal muscle atrophy via regulation of phosphatase tensin homolog. Hum Mol Genet, 26(10), 1821–1838. 10.1093/hmg/ddx085

12. Dai, N., Zhao, L., Wrighting, D., Kramer, D., Majithia, A., Wang, Y., Cracan, V., Borges-Rivera, D., Mootha, V. K., Nahrendorf, M., Thorburn, D. R., Minichiello, L., Altshuler, D., & Avruch, J. (2015). IGF2BP2/IMP2-Deficient mice resist obesity through enhanced translation of Ucp1 mRNA and Other mRNAs encoding mitochondrial proteins. Cell Metab, 21(4), 609–621. 10.1016/j.cmet.2015.03.006

13. Di Blasi, R., Marbiah, M. M., Siciliano, V., Polizzi, K., & Ceroni, F. (2021). A call for caution in analysing mammalian co-transfection experiments and implications of resource competition in data misinterpretation. Nat Commun, 12(1), 2545. 10.1038/s41467-021-22795-9

14. England, C. G., Ehlerding, E. B., & Cai, W. (2016). NanoLuc: A Small Luciferase Is Brightening Up the Field of Bioluminescence. Bioconjug Chem, 27(5), 1175–1187. 10.1021/acs.bioconjchem.6b00112

15. Enwerem, III, Elrod, N. D., Chang, C. T., Lin, A., Ji, P., Bohn, J. A., Levdansky, Y., Wagner, E. J., Valkov, E., & Goldstrohm, A. C. (2021). Human Pumilio proteins directly bind the CCR4- NOT deadenylase complex to regulate the transcriptome. RNA, 27(4), 445–464. 10.1261/rna.078436.120

16. Fang, Y., Liu, X., Liu, Y., & Xu, N. (2024). Insights into the Mode and Mechanism of Interactions Between RNA and RNA-Binding Proteins. Int J Mol Sci, 25(21). 10.3390/ijms252111337

17. Frei, T., Cella, F., Tedeschi, F., Gutierrez, J., Stan, G. B., Khammash, M., & Siciliano, V. (2020). Characterization and mitigation of gene expression burden in mammalian cells. Nat Commun, 11(1), 4641. 10.1038/s41467-020-18392-x

18. Gerstberger, S., Hafner, M., & Tuschl, T. (2014). A census of human RNA-binding proteins. Nat Rev Genet, 15(12), 829–845. 10.1038/nrg3813

19. Halbeisen, R. E., Galgano, A., Scherrer, T., & Gerber, A. P. (2008). Post-transcriptional gene regulation: from genome-wide studies to principles. Cell Mol Life Sci, 65(5), 798–813. 10.1007/s00018-007-7447-6

20. Hildebrandt, R. P., Moss, K. R., Janusz-Kaminska, A., Knudson, L. A., Denes, L. T., Saxena, T., Boggupalli, D. P., Li, Z., Lin, K., Bassell, G. J., & Wang, E. T. (2023). Muscleblind-like proteins use modular domains to localize RNAs by riding kinesins and docking to membranes. Nat Commun, 14(1), 3427. 10.1038/s41467-023-38923-6

21. Izumi, H., Imamura, T., Nagatani, G., Ise, T., Murakami, T., Uramoto, H., Torigoe, T., Ishiguchi, H., Yoshida, Y., Nomoto, M., Okamoto, T., Uchiumi, T., Kuwano, M., Funa, K., & Kohno, K. (2001). Y box-binding protein-1 binds preferentially to single-stranded nucleic acids and exhibits 3’-->5’ exonuclease activity. Nucleic Acids Res, 29(5), 1200–1207. 10.1093/nar/29.5.1200

22. Jones, R. D., Qian, Y., Siciliano, V., DiAndreth, B., Huh, J., Weiss, R., & Del Vecchio, D. (2020). An endoribonuclease-based feedforward controller for decoupling resource-limited genetic modules in mammalian cells. Nat Commun, 11(1), 5690. 10.1038/s41467-020-19126-9

23. Kaye, J. A., Rose, N. C., Goldsworthy, B., Goga, A., & L’Etoile, N. D. (2009). A 3’UTR pumilio-binding element directs translational activation in olfactory sensory neurons. Neuron, 61(1), 57–70. 10.1016/j.neuron.2008.11.012

24. Kleene, K. C. (2018). Y-box proteins combine versatile cold shock domains and arginine-rich motifs (ARMs) for pleiotropic functions in RNA biology. Biochem J, 475(17), 2769–2784. 10.1042/BCJ20170956

25. Kreiss, P., Cameron, B., Rangara, R., Mailhe, P., Aguerre-Charriol, O., Airiau, M., Scherman, D., Crouzet, J., & Pitard, B. (1999). Plasmid DNA size does not affect the physicochemical properties of lipoplexes but modulates gene transfer efficiency. Nucleic Acids Res, 27(19), 3792–3798. 10.1093/nar/27.19.3792

26. Lindquist, J. A., & Mertens, P. R. (2018). Cold shock proteins: from cellular mechanisms to pathophysiology and disease. Cell Commun Signal, 16(1), 63. 10.1186/s12964-018-0274-6

27. Lunde, B. M., Moore, C., & Varani, G. (2007). RNA-binding proteins: modular design for efficient function. Nat Rev Mol Cell Biol, 8(6), 479–490. 10.1038/nrm2178

28. Luo, E. C., Nathanson, J. L., Tan, F. E., Schwartz, J. L., Schmok, J. C., Shankar, A., Markmiller, S., Yee, B. A., Sathe, S., Pratt, G. A., Scaletta, D. B., Ha, Y., Hill, D. E., Aigner, S., & Yeo, G. W. (2020). Large-scale tethered function assays identify factors that regulate mRNA stability and translation. Nat Struct Mol Biol, 27(10), 989–1000. 10.1038/s41594-020-0477-6

29. Mordovkina, D., Lyabin, D. N., Smolin, E. A., Sogorina, E. M., Ovchinnikov, L. P., & Eliseeva, I. (2020). Y-Box Binding Proteins in mRNP Assembly, Translation, and Stability Control. Biomolecules, 10(4). 10.3390/biom10040591

30. Nie, M., Balda, M. S., & Matter, K. (2012). Stress- and Rho-activated ZO-1-associated nucleic acid binding protein binding to p21 mRNA mediates stabilization, translation, and cell survival. Proc Natl Acad Sci U S A, 109(27), 10897–10902. 10.1073/pnas.1118822109

31. Sahoo, P. K., Lee, S. J., Jaiswal, P. B., Alber, S., Kar, A. N., Miller-Randolph, S., Taylor, E. E., Smith, T., Singh, B., Ho, T. S., Urisman, A., Chand, S., Pena, E. A., Burlingame, A. L., Woolf, C. J., Fainzilber, M., English, A. W., & Twiss, J. L. (2018). Axonal G3BP1 stress granule protein limits axonal mRNA translation and nerve regeneration. Nat Commun, 9(1), 3358. 10.1038/s41467-018-05647-x

32. Sidibe, H., Dubinski, A., & Vande Velde, C. (2021). The multi-functional RNA-binding protein G3BP1 and its potential implication in neurodegenerative disease. J Neurochem, 157(4), 944–962. 10.1111/jnc.15280

33. Spitzer, J., Landthaler, M., & Tuschl, T. (2013). Rapid creation of stable mammalian cell lines for regulated expression of proteins using the Gateway(R) recombination cloning technology and Flp-In T-REx(R) lines. Methods Enzymol, 529, 99–124. 10.1016/B978-0-12-418687-3.00008-2

34. Sun, J., Yan, L., Shen, W., & Meng, A. (2018). Maternal Ybx1 safeguards zebrafish oocyte maturation and maternal-to-zygotic transition by repressing global translation. Development, 145(19). 10.1242/dev.166587

35. Tay, J., & Richter, J. D. (2001). Germ cell differentiation and synaptonemal complex formation are disrupted in CPEB knockout mice. Dev Cell, 1(2), 201–213. 10.1016/s1534-5807(01)00025-9

36. Uyhazi, K. E., Yang, Y., Liu, N., Qi, H., Huang, X. A., Mak, W., Weatherbee, S. D., de Prisco, N., Gennarino, V. A., Song, X., & Lin, H. (2020). Pumilio proteins utilize distinct regulatory mechanisms to achieve complementary functions required for pluripotency and embryogenesis. Proc Natl Acad Sci U S A, 117(14), 7851–7862. 10.1073/pnas.1916471117

37. Van Etten, J., Schagat, T. L., Hrit, J., Weidmann, C. A., Brumbaugh, J., Coon, J. J., & Goldstrohm, A. C. (2012). Human Pumilio proteins recruit multiple deadenylases to efficiently repress messenger RNAs. J Biol Chem, 287(43), 36370–36383. 10.1074/jbc.M112.373522

38. Van Nostrand, E. L., Freese, P., Pratt, G. A., Wang, X., Wei, X., Xiao, R., Blue, S. M., Chen, J. Y., Cody, N. A. L., Dominguez, D., Olson, S., Sundararaman, B., Zhan, L., Bazile, C., Bouvrette, L. P. B., Bergalet, J., Duff, M. O., Garcia, K. E., Gelboin-Burkhart, C., … Yeo, G. W. (2020). A large-scale binding and functional map of human RNA-binding proteins. Nature, 583(7818), 711–719. 10.1038/s41586-020-2077-3

39. Xu, D., Xu, S., Kyaw, A. M. M., Lim, Y. C., Chia, S. Y., Chee Siang, D. T., Alvarez-Dominguez, J. R., Chen, P., Leow, M. K., & Sun, L. (2017). RNA Binding Protein Ybx2 Regulates RNA Stability During Cold-Induced Brown Fat Activation. Diabetes, 66(12), 2987–3000. 10.2337/db17-0655

40. Zhang, Y., Cooke, A., Park, S., Dewey, C. N., Wickens, M., & Sheets, M. D. (2013). Bicaudal-C spatially controls translation of vertebrate maternal mRNAs. RNA, 19(11), 1575–1582. 10.1261/rna.041665.113

